# FibrilPaint: a class of amyloid-targeting peptides

**DOI:** 10.1101/2024.08.25.609586

**Authors:** Júlia Aragonès Pedrola, Françoise A. Dekker, Katrin Guttmann, Litske M. van Leeuwen, Shalini Singh, Guy Mayer, Tommaso Garfagnini, Assaf Friedler, Stefan G. D. Rüdiger

**Author notes:** These authors contributed equally (Júlia Aragonès Pedrola, Françoise A. Dekker).

## Abstract

Amyloid fibrils are a common pathological hallmark in multiple neurodegenerative diseases, yet molecular tools to selectively recognise and manipulate them remain scarce. We report FibrilPaints, a family of modular peptides designed for selective amyloid binding and adaptable chemical functionality. The degenerative amyloid-targeting unit of FibrilPaints, W_5_P_4_H_3_R_2_, has a high content of π-stacking and aromatic side chains. Systematic sequence variation, altering charge, termini, and residue order, revealed the importance of the composition of the amyloid-targeting unit for high-affinity binding across Tau and Huntingtin fibrils. Importantly, sequence changes outside this unit do not preclude fibril binding, which permits attachment of fluorophores or E3-recruiting motifs for targeted protein degradation. This work establishes FibrilPaint as a modular peptide system for the detection and modulation of amyloids.

Figure of content
The modular design of FibrilPaints enables systematic evaluation of their functionality by testing the binding capacity of each variant (FibrilPaintX) to distinct amyloid fibrils. Successful binding results in visible ‘painting’ of the fibrils, facilitating their detection and downstream research.

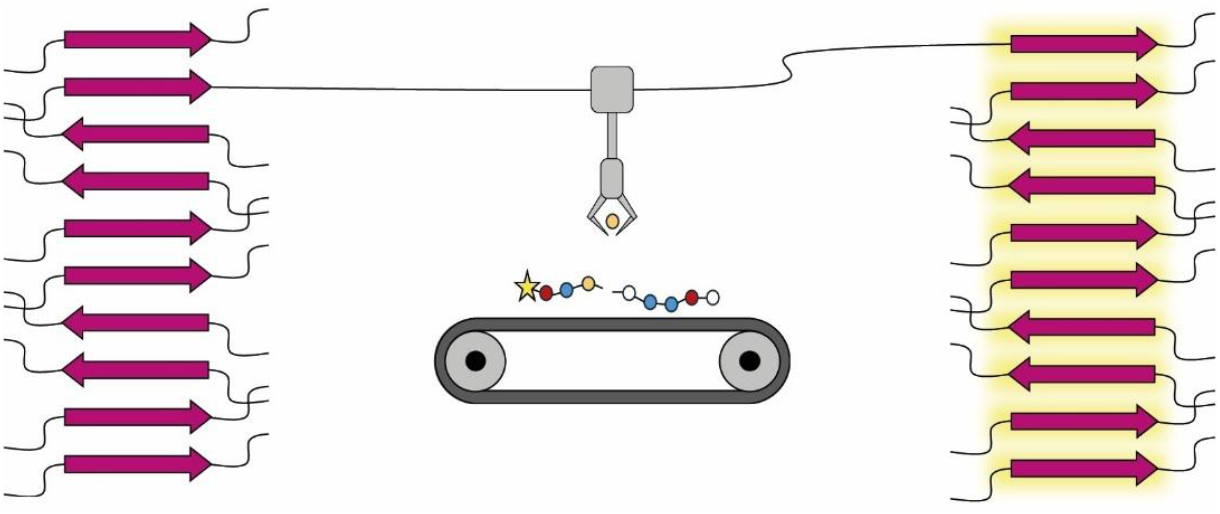

## Introduction

Neurodegenerative diseases such as Alzheimer’s (AD), Parkinson’s (PD), and Huntington’s Disease (HD) are defined by the accumulation of misfolded proteins into amyloid fibrils [1, 2]. These fibrillar assemblies share a cross-β structural motif that renders them highly stable and resistant to clearance [3, 4]. First amyloid targeting therapies have been approved, which delay the disease progression [5–8]. Although these therapies provide a valuable starting point, effects are still limited [9].

We therefore urgently need new compounds able to bind these amyloids. Peptides offer a versatile chemical platform, as their sequences can be tailored to exploit non-covalent interactions central to amyloid binding, including π–π stacking, cation–π interactions, and backbone hydrogen bonding [10, 11]. However, few peptide scaffolds allow systematic exploration of how sequence composition, charge, and stereochemistry influence fibril recognition. Developing modular peptide systems with predictable binding properties would therefore provide valuable tools for both diagnostic and therapeutic applications.

We recently introduced FibrilPaint1 (FP1), a 22-residue fluorescent peptide that binds a range of disease-associated amyloid fibrils, including Tau, Huntingtin (Htt), α-synuclein, and Amyloid-β, with affinity in the nanomolar range [12]. FP1 comprises an aromatic and arginine-rich core that drives amyloid interaction, a flexible linker, and a C-terminal EEVD motif capable of recruiting the E3 ligase CHIP [12, 13]. This design provides not only a binding module but also an adaptable sequence for conjugation or targeted protein degradation.

Here, we expand FP1 into a family of modular peptides; FibrilPaints; to define the molecular determinants of amyloid recognition. By systematically varying charge, sequence order, and termini, we identify derivatives that retain or lose binding to Tau and Htt fibrils. These studies reveal key features governing amyloid engagement and establish that the amyloid-targeting unit of FibrilPaints can be combined with entirely different C-terminal sequences. The resulting peptide class represents a tuneable chemical framework for developing amyloid-targeting probes and bifunctional molecules for degradation or imaging strategies.

## Results

### Design and development of a new class of amyloid-targeting peptides

FP1 binds multiple amyloid species, including Tau, Huntingtin (Htt), α-synuclein, and Aβ, with nanomolar affinity, yet it remained unclear which sequence features were essential for recognition and which could be modified for chemical versatility [12]. Now, we set out to expand this design into a broader family of amyloid-binding peptides. FP1 consists of three functional elements: (i) an aromatic and arginine-rich core sequence (PWWRRPWWPWHHPH) of which we hypothesise it mediates binding to amyloid β-sheets, (ii) a short glycine-serine linker (GSGS), and (iii) a C-terminal EEVD motif that recruits the E3 ligase CHIP for potential targeted degradation.

We designed a series of FP1 derivatives (FP2–FP13) differing in charge, residue order, length, and stereochemistry (Fig. 1, Table 1). Our goal was to define the sequence space compatible with amyloid recognition while introducing handles for future functionalisation. The design strategy included: (1) Charge variation, substituting acidic or basic residues to test electrostatic tolerance (FP5–FP7). (2) Sequence permutation, randomizing the aromatic–basic pattern to probe positional dependence (FP8). (3) Truncation, removing the linker or the EEVD motif to assess whether they contribute to binding (FP5–FP6). And (4) stereochemical inversion, replacing L-with D-amino acids to evaluate the role of backbone chirality (FP12–FP13). Each peptide retained a fluorescein (Fl) label at the N-terminus for follow-up analysis.

**Table 1.**
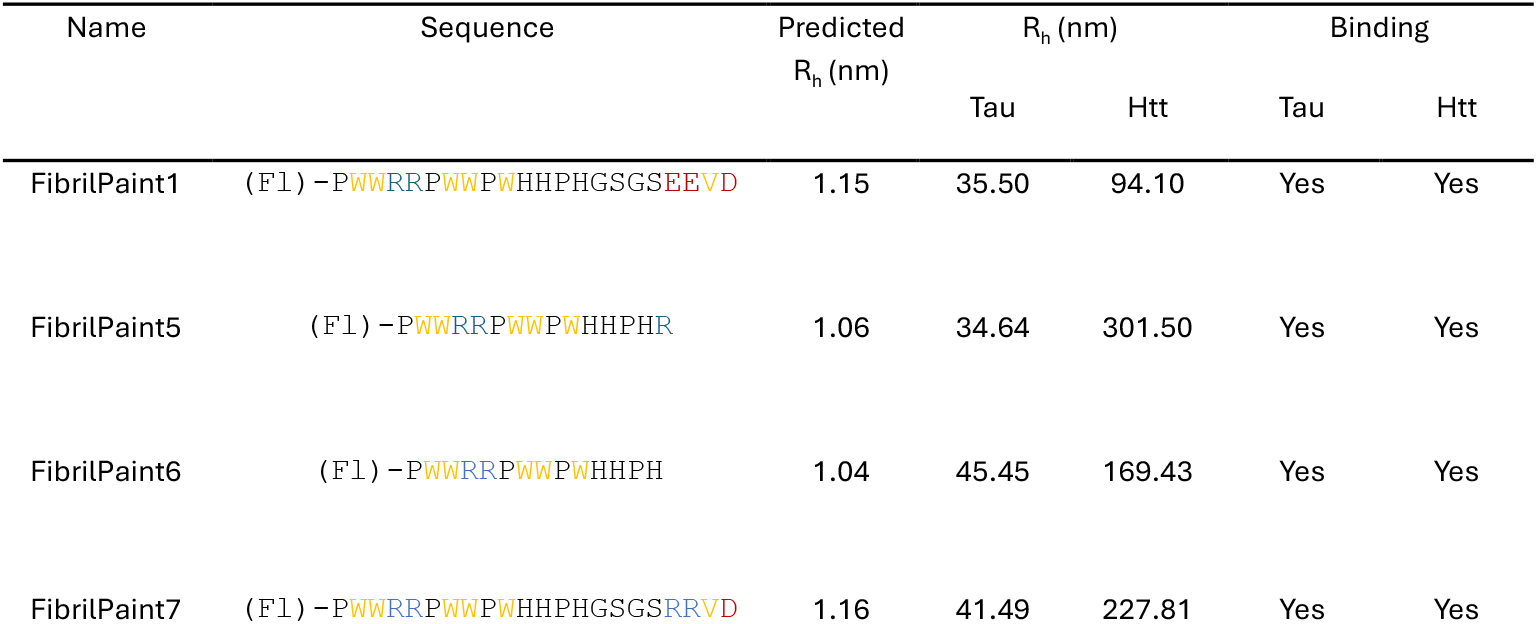
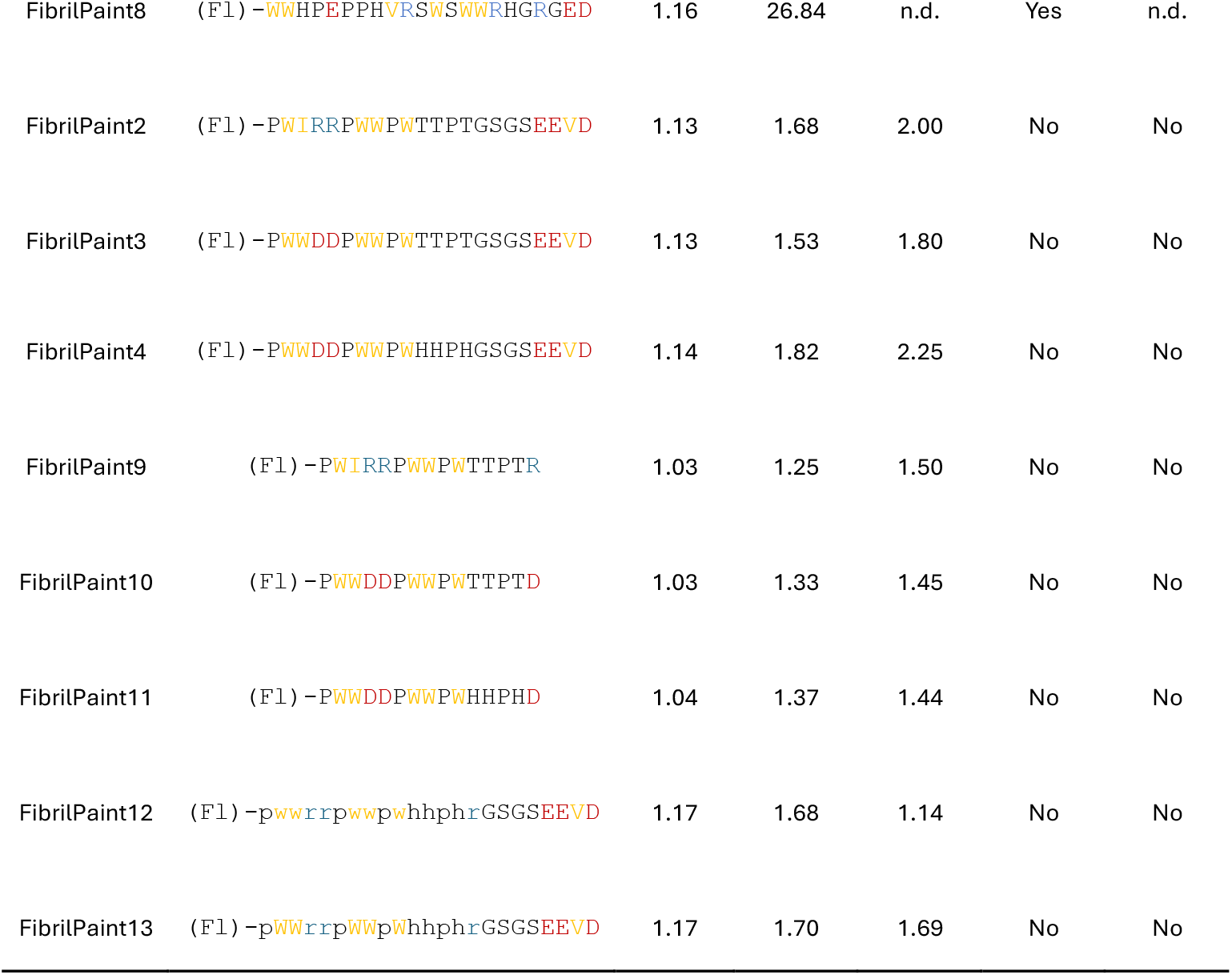
Properties of FibrilPaint peptides. All peptides were amidated at their C-termini. (Fl) indicates Fluorescein group. Amino acid residues are coloured as large hydrophobic/aromatic, yellow; negative charge, red; positive charge, blue. D-amino acid residues are referred as lower-case letters. Predicted Rh is the expected hydrodynamic radius based on the size of the peptide. The binding is assessed by the ability to measure the size of fibrils in a FIDA experiment

**Figure 1.**
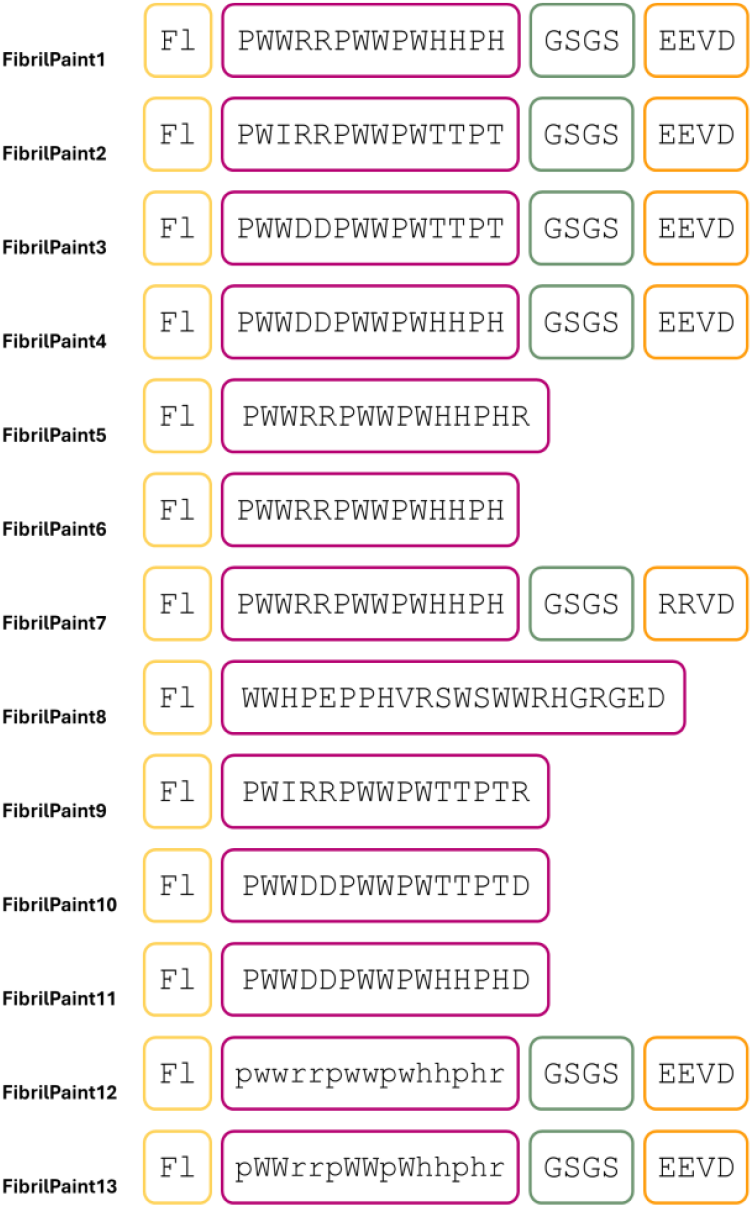
Modular design of FibrilPaint peptides. Schematic representation of the FibrilPaint architecture. Each peptide contains an N-terminal fluorescein tag (Fl, yellow), an amyloid-binding unit (purple) with sequence variations, and optional C-terminal motifs with linker (green) and motif for chemical functionalization or E3-ligase recruitment (orange).

### FibrilPaint Variants Retain Amyloid Recognition

Next, we aimed to determine the binding capacity of the FP peptides to amyloid fibrils, using Flow-Induced Dispersion Analysis (FIDA). FIDA is an established microfluidics and ensemble method, in which fluorescently labelled molecules disperse in solution [14, 15]. As a read-out it provides the Hydrodynamic Radius (R_h_) which is a measure for the cumulant radius of a biomolecular complex in solution. FP peptides alone, which are fluorescently labelled, have a predicted R_h_ ranging from 1.0 nm to 1.2 nm (Table 1). If FP binds to the fibrils, a larger complex is formed, and the measured size will increase. Thus, FIDA provides a direct measure of peptide binding.

To test the capacity of FP to bind amyloid fibrils, we used TauRD and HttEx1Q44 as model fibrils. Formation of Tau fibrils is associated to the development of AD [16], and Htt fibrilization is related to the progression of HD [17]. These two proteins differ in sequence and structure and only share the ability to form amyloid fibrils. We preformed TauRD and HttEx1Q44 fibrils, which resulted

We identified five active binders—FP1, FP5, FP6, FP7, and FP8—that consistently produced strong R_h_ increases after incubation with TauRD fibrils (Fig. 2). These peptides showed mean apparent radii around 37 nm. We repeated the experiment with HttEx1Q44 fibrils, where R_h_ values increased to 188 nm (Fig. 3). The consistent fibril-dependent size shifts confirm specific association of these peptides with amyloid aggregates, independent of minor sequence variations. The fibril size variation within the measurement using different FPs is minimal and does not depend on the FP. Given that the aggregation process is a seeded event, it is expected to observe some variation in the measured Rh. Together, these results indicate that FP1, FP5, FP6, FP7 and FP8 are effective fibril binders.

**Figure 2.**
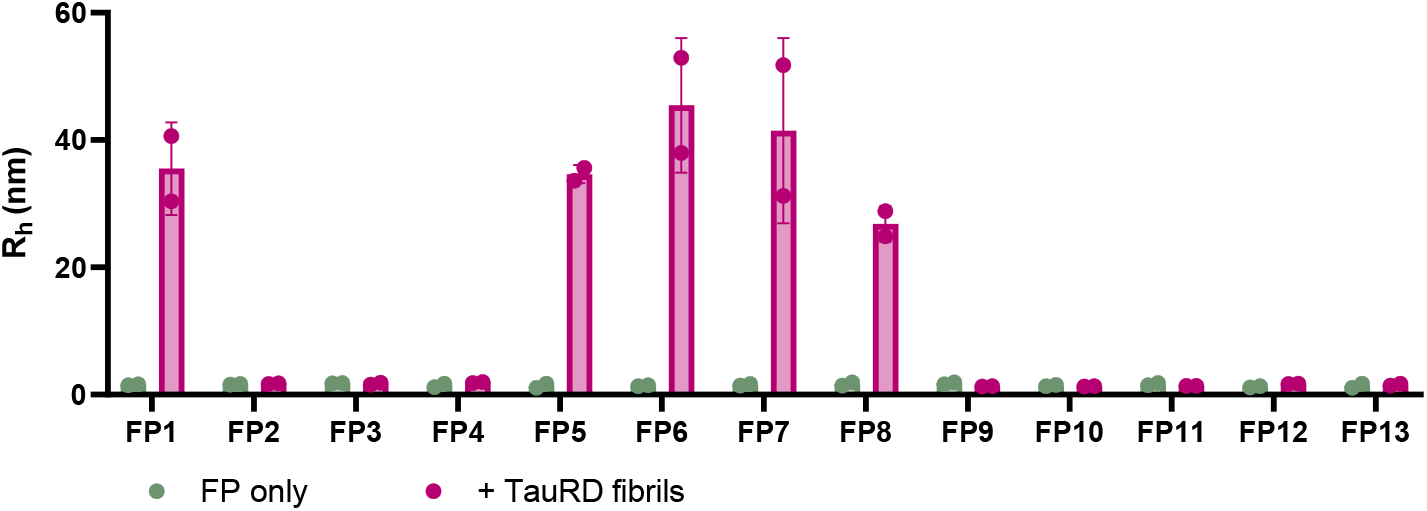
Hydrodynamic radius (R_h_) ofFibrilPaint variants incubated with TauRD fibrils. R_h_ measurement with Flow-Induced Dispersion Analysis (FIDA) of fluorescently labelled FibrilPaint peptides after 15 minutes incubation with preformed TauRD fibrils (pink) or without (green).

**Figure 3.**
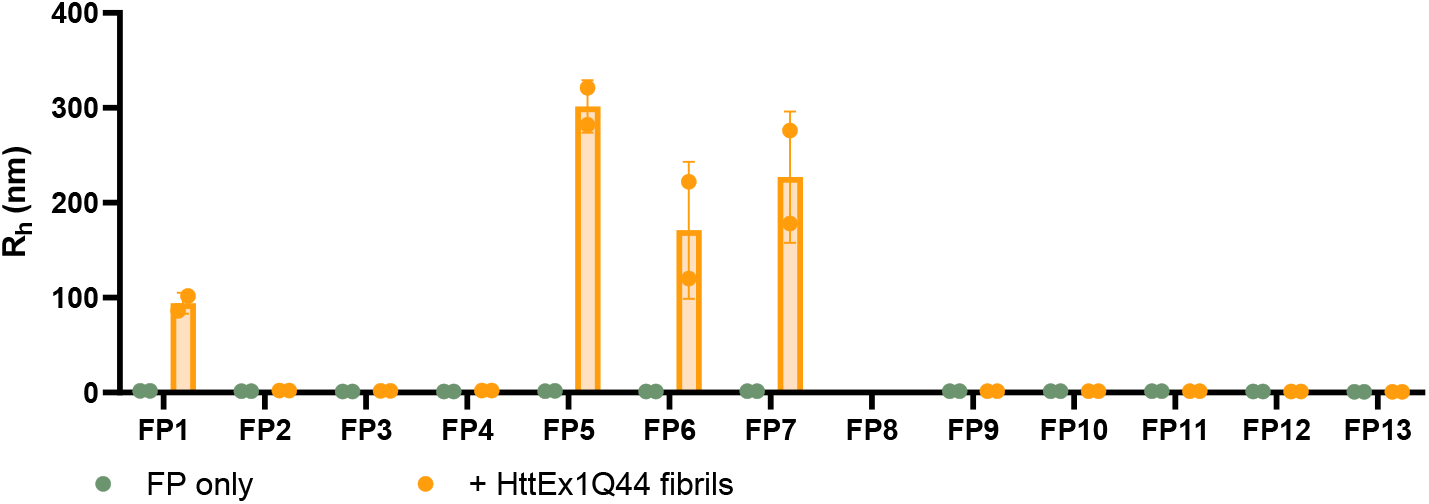
Hydrodynamic radius (R_h_) of FibrilPaint variants incubated with HttEx1Q44 fibrils. R_h_ measurement with Flow-Induced Dispersion Analysis (FIDA) of fluorescently labelled FibrilPaint peptides after 15 minutes incubation with preformed Huntingtin fibrils (orange) or without (green).

By contrast, peptides FP2–FP4 and FP9–FP13 showed no measurable binding (Fig. 2, 3). indicating that certain substitutions, particularly loss of arginine or histidine residues, or inversion to D-chirality, disrupt the molecular interactions required for fibril engagement.

### The sequence of Fibril Paint peptides can be modified while maintaining specific recognition features of the of amyloid-targeting unit

The identification of four new FibrilPaint peptides binding amyloid fibrils (FP5, FP6, PF7 and FP8) demonstrates that the sequence of FP1 can be varied while maintaining specific recognition features of the of amyloid-targeting unit. First, an overall change in charge does not compromise binding ability. FP1 has a net charge of −1, and FP5, FP6, FP7 and FP8 have a net charge of +3, +2, +2, and 0, respectively. FP5, FP6 lack the GSGS and the EEVD motif, and in FP7 EEVD has been replaced by RRVD, indicating that those motifs are not required for detection of amyloid fibrils. In FP8, the order of the amino acid sequence is altered, indicating a potential to vary the sequential order of the residues.

FP9, FP10, FP11, FP12 and FP13 lost ability to bind fibrils, suggesting that some components of the sequence are important to preserve the binding capacity. Residues R and H seem to play an essential role in binding, since the loss of these residues compromised the binding capacity of FP9, FP10 and FP11. Remarkably, the backbone conformation is important, considering that FP12 and FP13, which have D-amino acids, do not detect amyloid fibrils in FIDA measurements.

Together, these data define a new family of amyloid-targeting FibrilPaint peptides that preserve binding despite variations in charge, linker, or sequence order. The aromatic–basic core thus represents the essential recognition motif, while other regions of the peptide can be freely adapted for chemical or functional modification.

## Discussion

In this study, we establish FibrilPaints as a modular class of amyloid-binding peptides. Building on the FP1 scaffold, systematic sequence variation revealed which molecular features are essential for amyloid recognition and which can be freely modified [18]. Four new derivatives (FP5, FP6, FP7, and FP8) retained strong binding to Tau and Huntingtin fibrils, despite alterations in overall charge, termini, and sequence order. This highlights the importance of the sequence features of the amyloid targeting unit of FibrilPaint peptides, which provides a versatile framework for engineering next-generation amyloid-targeting molecules.

The sequence composition of the amyloid-binding unit of the FibrilPaint ancestor FP1 is degenerate, as it contains only four different residues on a length of 14 amino acids: W_5_P_4_H_3_R_2_. All 14 residues have polar groups. Ten of them contain delocalised π-systems, eight of which are aromatic. This is consistent with π–π and cation–π interactions that mediate the association of peptides with β-sheet– rich fibrils [10, 19–21]. Our defined sequence alterations show that these are critical features for the amyloid-binding properties of FibrilPaints. Removal of an aromatic residue (FP2 and FP9) resulted in losing fibril binding, so did an exchange of both Arg residues with Asp. The overall amino acid composition of FibrilPaints is more important than the sequence order, as FP8 still binds to Tau fibrils. This is in agreement with recent reports that aggregation inhibition depends on physicochemical composition rather than primary sequence order [11, 22, 23]. In contrast, inversion of chirality (D-amino acid substitution) prevented binding, underscoring a stereochemical dependence of peptide-fibril complementarity.

Important for potential adaption of FibrilPaints to various cellular processes is that the sequence outside of the amyloid-binding unit can be varied. It can be deleted altogether (FP6) and its charge can be inverted without compromising fibril binding (FP5 and FP7). Beyond elucidating binding determinants, the chemical modularity of FibrilPaints makes them attractive building blocks for bifunctional applications. The EEVD terminus of FP1 recruits the E3 ligase CHIP, enabling the concept of targeted protein degradation of amyloid fibrils [13]. The modular design allows this recruiting element to be replaced by alternative chemical handles or degradation motifs, such as autophagy-targeting tags, expanding the scope from proteasomal to lysosomal clearance [24–27]. In this sense, FibrilPaints bridge molecular recognition and functional intervention, similar to PROTAC and AUTAC strategies, but adapted to supramolecular protein assemblies. The present work demonstrates that both halves of the FibrilPaint design; the amyloid-binding unit and the effector module; can be independently optimised, providing a rational route to multifunctional peptide therapeutics.

In summary, FibrilPaints present a peptide system that allows systematic chemical tailoring for the selective recognition and manipulation of amyloid fibrils. Their modularity, structural robustness, and compatibility with diverse functional groups make them promising scaffolds for new chemical biology tools targeting protein aggregation, and maybe later guide the development of new imaging probes to visualise amyloids in patients, or degrader molecules effectively targeting only amyloids. By bridging molecular design with biological relevance, this work lays the foundation for translating amyloid recognition into therapeutic and diagnostic applications.

## Conclusions

This work establishes FibrilPaints as a modular and chemically versatile class of peptides for the selective recognition of amyloid fibrils. Through systematic sequence variation, we identified key structural determinants, aromatic and basic residues with π-stacking ability, that govern fibril binding, while demonstrating that charge, termini, and sequence order can be flexibly tuned without loss of activity. The resulting scaffolds provide a foundation for chemical diversification, enabling conjugation to dyes, ligase-recruiting motifs, or other functional groups. By establishing an amyloid recognition motif with chemical adaptability, FibrilPaints offer a generalisable strategy for designing peptide-based tools to detect, modulate, or eliminate amyloid aggregates in neurodegenerative disease.

## Author contributions

Conception JAP, FAD, TG, AF, SGDR

Design of the work JAP, FAD, SGDR

Acquisition JAP, FAD, KG, LML, SS, TG, GM

Analysis JAP, FAD, KG, LML, SGDR

Interpretation of data JAP, FAD, KG, LML, SGDR

Writing original draft JAP

Revision JAP, FAD, TG, AF, SGDR

## Data availability

Data for this article is provided as additional information document.

## Acknowledgements

SGDR was supported by grants of the Campaign Team Huntington and Alzheimer Nederland (No. WE.03-2019-03) and a ZonMW TOP grant (No. 91215084) and is a principal investigators of the Gravitation Consortium “FLOW” (024.006.036), funded by the Dutch Ministry of Education, Culture, and Science (OCW). FAD and SGDR were supported by NWO TakeOff1 (22073) and BioTech Booster (BIOB25014). AF thanks The Minerva Center for Bio-Hybrid complex systems and the Saerree K. and Louis P. Fiedler Chair in Chemistry. Measurements on the FIDA1, CLARIOstar® Plus and Monolith were done at the Protein Research Centre of Utrecht University.

## Methods

### Peptide synthesis and purification

The peptides were synthesized on a Rink amide resin using a Liberty Blue Microwave-Assisted Peptide Synthesizer (CEM) with standard Fmoc chemistry and Oxyma/DIC as coupling reagents. The peptide concentrations were measured by UV spectroscopy. The peptides were labelled with 5(6)-carboxyfluorescein at their N’ termini. The peptides were cleaved from the resin with a mixture of 95 % (v/v) trifluoroacetic acid (TFA), 2.5% (v/v) triisopropylsilane (TIS), 2.5 % (v/v) triple distilled water (TDW) agitating vigorously for 3 hours at room temperature. The volume was decreased by N2 flux, and the peptides precipitated by addition of 4 volumes of diethylether at −20 °C. The peptides were sedimented at −20 °C for 30 minutes, then centrifuged and the diethylether discarded. The peptides were washed three times with diethylether and dried by gentle N2 flux. The solid was dissolved in 1:2 volume ratio of acetonitrile (ANC):TDW, frozen in liquid Nitrogen and lyophilized. The peptides were purified on a WATERS HPLC using a reverse-phase C18 preparative column with a gradient ACN/TDW. The identity and purity of the peptides was verified by ESI mass spectrometry and Merck Hitachi analytical HPLC using a reverse-phase C8 analytical column.

### Expression and purification of TauRD

We produced N-terminally FLAG-tagged (DYKDDDDK) human TauRD (Q244-E372, with pro-aggregation mutation ΔK280) in E. Coli BL21 Rosetta 2 (Novagen), with an additional removable N-terminal His6-Smt-tag (MGHHHHHHGSDSEVNQEAKPEVKPEVKPETHINLKVSDGSSEIFFKIKKTTPLRRLMEAFAKRQGKEMDSLRFLYDGIRIQADQTPEDLDMEDNDIIEAHREQIGG). Expression was induced at OD600 0.8 by addition of 0.15 mM IPTG and incubation at 18 °C overnight. Cells were harvested by centrifugation, resuspended in 25 mM HEPES-KOH pH=8.5, 50 mM KCl, flash frozen in liquid nitrogen, and kept at – 80 °C until further usage. Pellets were thawed at 37 °C, followed by the addition of ½ tablet/50 ml EDTA-free Protease Inhibitor and 5 mM β-mercaptoethanol. Cells were disrupted using an EmulsiFlex-C5 cell disruptor, and lysate was cleared by centrifugation. Supernatant was filtered using a 0.22 μm polypropylene filtered and purified with an ÄKTA purifier chromatography System. Sample was loaded into a POROS 20MC affinity purification column with 50 mM HEPES-KOH pH 8.5, 50 mM KCl, eluted with a linear gradient 0-100%, 5 CV of 0.5 M imidazole. Fractions of interest were collected, concentrated to 2.5 ml using a buffer concentration column (vivaspin, MWCO 10 KDa), and desalted using PD-10 desalting column to HEPES pH 8.5, ½ tablet/50 ml Complete protease inhibitor, 5 mM β-mercaptoethanol. The His6-Smt-tag was removed by treating the sample with Ulp1, 4 oC, shaking, overnight. The next day, sample was loaded into POROS 20HS column with HEPES pH 8.5, eluted with 0-100% linear gradient, 12 CV of 1M KCl. Fractions of interest were collected and loaded into a Superdex 26/60, 200 pg size exclusion column with 25 mM HEPES-KOH pH 7.5, 75 mM NaCl, 75 mM KCl. Fractions of interest were concentrated using a concentrator (vivaspin, MWCO 5 kKDda) to desired concentration. Protein concentration was measured using a NanoDrop™ OneC UV/Vis spectrophotometer and purity was assessed by SDS-PAGE. Protein was aliquoted and stored at – 80 °C.

### Expression and purification of HttEx1Q44

We produced HttEx1Q44 with in E. Coli BL21 Rosetta 2 (Novagen), with an additional N-terminal MBP-and C-terminal His6-tag (MKIEEGKLVIWINGDKGYNGLAEVGKKFEKDTGIKVTVEHPDKLEEKFPQVAATGDGPDIIFWAHDRFGGYAQSGLLAEITPDKAFQDKLYPFTWDAVRYNGKLIAYPIAVEALSLIYNKDLLPNPPKTWEEIPALDKELKAKGKSALMFNLQEPYFTWPLIAADGGYAFKYENGKYDIKDVGVDNAGAKAGLTFLVDLIKNKHMNADTDYSIAEAAFNKGETAMTINGPWAWSNIDTSKVNYGVTVLPTFKGQPSKPFVGVLSAGINAASPNKELAKEFLENYLLTDEGLEAVNKDKPLGAVALKSYEEELAKDPRIAATMENAQKGEIMPNIPQMSAFWYAVRTAVINAASGRQTVDAALAAAQTNAAAASEFSSNNNNNNNNNNLGIEGRMATLEKLMKAFESLKSFQQQQQQQQQQQQQQQQQQQQQQQQQQQQQQQQQQQQQQQQQQQQPPPPPPPPPPPQLPQPPPQAQPLLPQPQPPPPPPPPPPGPAVAEEPLHRPSGSHHHHHH). We induced expression at an OD600 of 0.8 by adding 0.15 mM IPTG and incubating at 18 °C overnight. Cells were harvested by centrifugation and resuspended in 50 mM HEPES-KOH pH=8.5, 50 mM KCl with ½ tablet/50 ml EDTA-free Protease Inhibitor and 5 mM β-mercaptoethanol. Cells were disrupted using an EmulsiFlex-C5 cell disruptor, and lysate was cleared by centrifugation.

Supernatant was filtered using a 0.22 μm polypropylene filtered and purified with an ÄKTA purifier chromatography System. Sample was loaded into a POROS 20MC affinity purification column with 50 mM HEPES-KOH pH 8.5, 100 mM KCl, eluted with a linear gradient 0-100%, 10 CV of 0.5 M imidazole, 30 mM NaCl. Fractions of interest were collected, buffer exchanged to 50 mM HEPES 150 mM NaCl and concentrated to desired concentration using a Vivaspin column (vivaspin, MWCO 30 KDa).

Protein concentration was measured using a NanoDrop™ OneC UV/Vis spectrophotometer and purity was assessed by SDS-PAGE. Protein was aliquoted and stored at – 80 °C.

### TauRD fibril formation

Aggregation of 20 μM TauRD in 25 mM HEPES-KOH pH 7.4, 75 mM KCl, 75 mM NaCl, ½ tablet/50 ml Protease Inhibitor, was induced by the addition 5 μM of heparin low molecular weight and incubated at 37 °C and 600 rpm in an EppendorfTM Thermomixer® during 20 hours. Fibrils were made independently, per measurement.

### HttEx1Q44 fibril formation

Aggregation of 20 μM HttEx1Q44 in 25 mM HEPES-KOH pH 7.4, 75 mM KCl, 75 mM NaCl, ½ tablet/50 ml Protease Inhibitor, was induced by the cleavage of C-terminal MBP-tag with Factor Xa and incubated at 37 °C and 600 rpm in an EppendorfTM Thermomixer® during 5 hours. Fibrils were made independently, per measurement.

### Size measurement with Flow Induced Dispersion Analysis

200 nM of Fibril Paint was measured in presence or absence of 2 μM preformed TauRD or HttEx1Q44 fibrils in 25 mM HEPES-KOH pH 7.5, 75 mM KCl, 75 mM NaCl, 0.5% pluronic (for TauRD) or 50 mM HEPES-KOH pH 7.5, 150 mM KCl, 0.5% pluronic (for HttExQ44). Size measurements were performed in a FIDA1 instrument with a 480 nm excitation source, using a capillary dissociation (Capdis) approach. In this, the capillary is equilibrated with buffer followed by injection of the sample. Tray 1 was maintained at 37 °C and tray 2 and capillary chamber at 25 °C

**Table 1.** Experimental parameters for analysis of proteins up to Rh of 10 nm.

| Tray | Vial | Pressure<br>(mbar) | Time<br>(s) | Outlet | Measure | Comments |
| --- | --- | --- | --- | --- | --- | --- |
| 2 | 1 | 3500 | 45 | Variable | No | 1 M NaOH |
| 2 | 2 | 3500 | 75 | Variable | No | Buffer<br>equilibration |
| 1 | Analyte | 3500 | 20 | Variable | No | Buffer |
| 1 | Indicator | 50 | 10 | Variable | No | Sample<br>application |
| 1 | Analyte | 400 | 180 | Variable | Yes | Buffer |

Tray 1 was maintained at 37 oC and tray 2 and capillary chamber at 25 oC

**Table 2.** Experimental parameters for analysis of proteins up to Rh of 75 nm.

| Tray | Vial | Pressure<br>(mbar) | Time<br>(s) | Outlet | Measure | Comments |
| --- | --- | --- | --- | --- | --- | --- |
| 2 | 1 | 3500 | 45 | Variable | No | 1 M NaOH |
| 2 | 2 | 3500 | 75 | Variable | No | Buffer<br>equilibration |
| 1 | Analyte | 3500 | 20 | Variable | No | Buffer |
| 1 | Indicator | 50 | 10 | Variable | No | Sample<br>application |
| 1 | Analyte | 75 | 1800 | Variable | Yes | Buffer |

